# Assessment of Heterogeneity of Cytochrome P450 Activity in Cancer-Cell Population by Cytometry of Reaction Rate Constant is Robust to Variation in Substrate Concentration

**DOI:** 10.1101/2020.10.15.341289

**Authors:** Mariana Bleker de Oliveira, Vasilij Koshkin, Christopher G. R. Perry, Sergey N. Krylov

**Affiliations:** Centre for Research on Biomolecular Interactions, York University, Toronto M3J 1P3, Canada

## Abstract

Enzymes of the cytochrome P450 (CYP) family catalyze the metabolism of chemotherapeutic agents and are among the key players in primary and acquired chemoresistance of cancer. The activity of CYP is heterogeneous in tumor tissues, and the quantitative characteristics of this heterogeneity can be used to predict chemoresistance. Cytometry of reaction rate constant (CRRC) is a kinetic approach to assess cell population heterogeneity by measuring rates of processes at the single-cell level *via* time-lapse imaging. CRRC was shown to be an accurate and robust method for assessing the heterogeneity of drug-extrusion activity catalyzed by ABC transporters, which are also key players in cancer chemoresistance. We hypothesized that CRRC is also a reliable method for assessing the heterogeneity of CYP activity. Here, we evaluated the robustness of assessing the heterogeneity of CYP activity by CRRC with respect to controlled variation in the concentration of a CYP substrate by comparing CRRC with non-kinetic approaches. We found that changing the substrate concentration by 20% resulted only in minimal changes in the position, width, and asymmetry of the peak in the CRRC histogram, while these parameters varied greatly in the non-kinetic histograms. Moreover, the Kolmogorov-Smirnov statistical test showed that the distribution of the cell population in CRRC histograms was not significantly different; the result was opposite for non-kinetic histograms. In conclusion, we were able to demonstrate the robustness of CRRC with respect to changes in substrate concentration when evaluating CYP activity at the single-cell level.

## Introduction

Cytometry of reaction rate constant (CRRC) is a kinetic approach to identify cell population heterogeneity by measuring rates of cellular processes at the single-cell level *via* time-lapse imaging [1]. Identification of heterogeneous populations of cells is critical in predicting cancer chemoresistance because resistance to primary chemotherapy is primarily caused by a small drug-resistant subpopulation of tumor cells with activated cellular process of drug extrusion by ATP-binding cassette (ABC) transporters and drug degradation by metabolic enzymes [2]. The relative size of the drug-resistant subpopulation has been shown to correlate with clinical chemoresistance; hence, it is considered to be a marker of chemoresistance [3, 4].

Finding a predictor for chemoresistance based on the size of the drug-resistant subpopulation requires accurate determination of this size via monitoring the activity of the aforementioned processes at the single-cell level [5]. We have demonstrated that CRRC can characterize cell population heterogeneity based on the activity of ABC transporters, and that CRRC measurements are robust to changes in substrate concentration [1]. This robustness is paramount, since volumes of cell suspensions and substrates can vary up to 20% in repeated experimental settings, due to systematic (calibration of instruments) or random (incorrect pipetting) errors [6].

After extensively studying ABC transporters, we now aim to demonstrate that CRRC can also be robust to substrate-concentration variation in assessing cell-population heterogeneity for activities of other processes, specifically, drug degradation by metabolic enzymes. Cytochrome P450 (CYP) enzymes catalyze drug metabolism; the activity of this metabolism correlates with chemoresistance in several types of cancer [7–10]. Therefore, the goal of this study is to evaluate the robustness of CRRC with respect to CYP substrate-concentration variation.

## Materials and Methods

### Cell culture

HepG2 cells were donated by Dr. Christopher Perry (School of Kinesiology & Health Science, York University, ON, Canada). Cells were grown in EMEM media, supplemented with 100 IU/mL penicillin, 100 μg/mL streptomycin, and 10% fetal bovine serum, at 37 °C, in a humidified atmosphere of 5% CO_2_.

### Image acquisition

To assess CYP activity, we followed o-dealkylation of pentoxyresorufin (a fluorogenic CYP substrate) forming resorufin (a fluorescent product) over time with a Leica DMi8 microscope. Pentoxyresorufin is lipophilic and was diluted in DMSO. Cells were plated in a 35 mm dish, at 60–70% confluence, in HBSS medium. Probenecid (1 mM), an organic-anion transporter inhibitor, was added to prevent extrusion of the product and dicoumarol (25 μM), an inhibitor of the enzyme NAD(P)H Quinone Dehydrogenase 1, was added to inhibit further transformation of the product (resorufin) into non-fluorescent dihydroresorufin. Time-lapse fluorescence imaging of the cells was started, and an image of the field of view was collected every minute for 20 min. Cells were maintained at room temperature during time-lapse imaging. After registration of background fluorescence for 3 min, the substrate, pentoxyresorufin, was added to the cells at concentrations of either 5 μM or 6 μM. Snapshots were taken using a 10× lens and images were taken with the high-sensitivity camera mode. The identification of individual cells was performed with propidium iodide and saponin staining after image acquisition was over.

### Extraction and Analysis of Kinetic Traces

Values of fluorescence intensity were obtained from each cell using ImageJ software for 1,235 cells for each substrate concentration. OriginPro software was used to obtain kinetic traces of each cell and for the following kinetic and non-kinetic evaluations. For the kinetic assessment with CRRC, the value of the first order rate constant *k*_CYP_ was determined from the exponential fit of each kinetic trace. Two non-kinetic approaches were utilized: initial rate and average rate. The initial rate was calculated as the fluorescence intensity increase one minute after substrate addition:

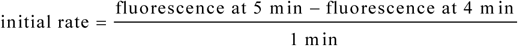

The average rate was calculated as the averaged fluorescence intensity increase over a 10-min period after substrate addition:

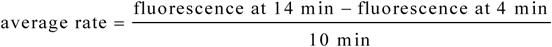

Note, that while *k*_CYP_, initial rate, and average rate have units of min^−1^, their absolute values differ significantly and are not to be compared.

### Cell Population Analysis

Three types of histograms were plotted in OriginPro software using the automatic binning mode: 1) number of cells vs *k*_CYP_, 2) number of cells vs initial rate, and 3) number of cells vs average rate. The histograms were characterized by three quantitative parameters, namely: median values, interquartile range, and skewness. Distributions in the histograms were compared with the Kolmogorov-Smirnov test, considering *p* < 0.05 as a criterion of statistical significance.

## Results and Discussion

Histograms for CRRC, initial rate, and average rate are shown in Figure 1. To evaluate the robustness, we varied he concentration of CYP substrate by 20%: 5 μM and 6 μM. Kinetic and non-kinetic histograms are unimodal, suggesting one single population of cells, which was expected for a pure single cell line.

**Figure 1.**
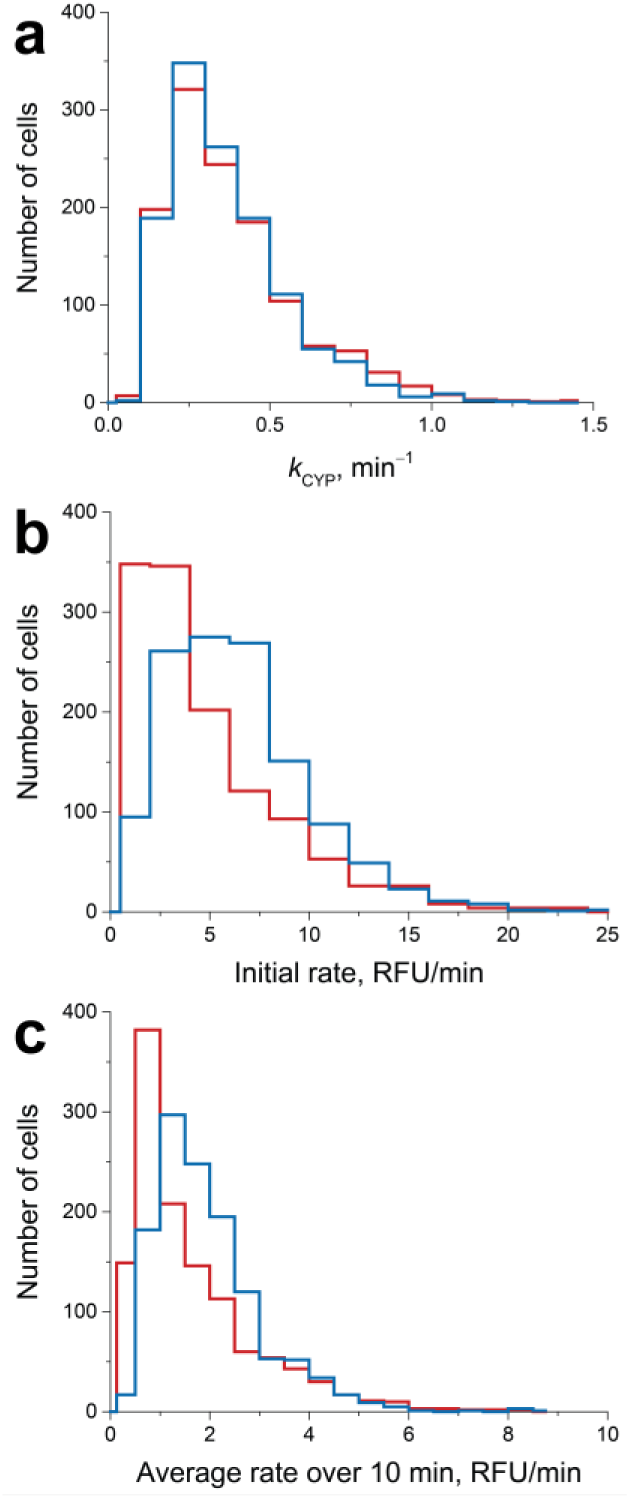
Kinetic (a) and non-kinetic (b and c) histograms for CYP activity obtained from single-cell analysis when varying substrate concentration by 20%: 5 μM (red line) and 6 μM (blue line).

Histograms were characterized by three values: median value of *x* (peak position), interquartile range (peak width), and skewness (peak asymmetry). Varying the substrate concentration did not cause significant changes in the kinetic histograms. The peak position remained the same as the median value of *k*_CYP_, was 0.33 min^−1^ for both 5 μM and 6 μM substrate concentrations (Figure 1a). The values of parameters characterizing cell-population heterogeneity for the two substrate concentrations can be seen in Table 1. Interquartile range and skewness varied by 11.7% and 16.9%, respectively.

**Table 1.** Quantitative characteristics of cell-population heterogeneity (median value of *x*, interquartile range, and skewness) at 5 and 6 µM substrate concentrations for the kinetic histogram (*k*_CYP_) and two non-kinetic histograms (initial rate and average rate).

| Parameter | Values for 5 $\mu\text{M}$ | Values for 6 $\mu\text{M}$ | Variation |
| --- | --- | --- | --- |
| **$k_{\text{CYP}}$** |  |  |  |
| Median $x$ | $0.33 \text{ min}^{-1}$ | $0.33 \text{ min}^{-1}$ | 0% |
| Interquartile range | $0.25 \text{ min}^{-1}$ | $0.22 \text{ min}^{-1}$ | 11.7% |
| Skewness | 1.96 | 1.62 | 16.9% |
| **Initial rate** |  |  |  |
| Median $x$ | 3.46 RFU/min | 5.88 RFU/min | 41.1% |
| Interquartile range | 4.70 RFU/min | 4.65 RFU/min | 1% |
| Skewness | 1.54 | 1.10 | 28.4% |
| **Average rate** |  |  |  |
| Median $x$ | 1.17 RFU/min | 1.71 RFU/min | 31.7% |
| Interquartile range | 1.47 RFU/min | 1.29 RFU/min | 12.6% |
| Skewness | 1.68 | 1.56 | 7.4% |

For initial and average rates, changes in peak position for different substrate concentrations were substantial. Median values of *x* for initial rate were 3.46 for 5 μM and 5.88 for 6 μM (Figure 1b), which corresponds to a variation of 41.1%. Variation of interquartile range was 1% and that of skewness was 28.4%. For average rate, median values of *x* were 1.17 for 5 μM and 1.71 for 6 μM (Figure 1c), resulting in a variation of 31.7%. Interquartile range varied by 12.6% and skewness by 7.4%. Moreover, in the kinetic histograms, distributions were not significantly different (*p* < 0.05), while distributions in the non-kinetic histograms were statistically different (*p* < 0.05).

Robust histograms should be superimposable and peak positions should be similar, with a negligible range of variation. This criterion was satisfied for kinetic histograms with *k*_CYP_ being the measure of CYP activity. In contrast, the non-kinetic histograms, with the initial and average rates being the measure of CYP activity, were non-robust; that is, 20% of variation in substrate concentration resulted in significant changes in peak position. Only minimal variation for peak position was observed in the kinetic assessment using CRRC, with the same median *k*_CYP_ for both substrate concentrations, while variation of peak position was over 30% for both non-kinetic approaches. Although variations of peak widths (interquartile range) and asymmetry (skewness) were not the lowest for CRRC, they were still small in absolute values (below 20%). Moreover, the kinetic histogram was the only one that had a cell population distribution that was not statistically different. Thus, CRRC is robust to variations in substrate concentration, while the non-kinetic approaches are not.

Our entire CRRC development to this point was carried out with a single drug-resistance process: drug extrusion from cells by ABC transporters [1, 11, 12]. These previous studies proved that the accuracy and robustness of CRRC greatly exceed those of the currently-used non-kinetic approach of assessing ABC activity based on measurements of ABC substrate retention, demonstrating the expected high utility of this technique.

The goal of the present study was to evaluate another important mechanism of chemoresistance in cancer cells: drug metabolism. There are only a few studies analyzing kinetics in drug metabolizing enzymes, at the single cell level [13–15], and they were not followed up on nor were their conclusions applied clinically. As for CYP, more recent studies of CYP activity in single cells have focused their analysis on specific rates of fluorescence change rather than chemical reaction parameters, such as rate constant [16–18]. Those specific rates, unlike rate constants, strongly depend on particular assay conditions and are unsuitable for broad comparative analysis.

## Conclusions

In conclusion, we were able to show that CRRC histograms are robust to variation in substrate concentration when evaluating CYP activity at the single-cell level. Non-kinetic histograms are non-robust to this variation. In the future, we aim at evaluating the accuracy of CRRC with respect to CYP activity and its ability to characterize distinct cell populations with different levels of CYP activity.

